# Shared lipidome and proteome signatures of frontotemporal lobar degeneration and Alzheimer’s disease

**DOI:** 10.64898/2026.07.11.737778

**Authors:** Yohannes A. Ambaw, Alissa L. Nana, Zhuoning Li, Shubham Singh, Mara Monetti, Bruce L. Miller, Salvatore E. Spina, Lea T. Grinberg, William W. Seeley, Tobias C. Walther, Robert V. Farese

**Author notes:** These authors contributed equally to this work (Tobias C. Walther, Robert V. Farese).

## Abstract

Frontotemporal lobar degeneration (FTLD) and Alzheimer’s disease (AD) differ in their clinical features and genetic etiologies but share progressive cognitive decline. Emerging evidence implicates lipid dysregulation in neurodegeneration, but its extent across FTLD subtypes and how it compares to AD are unclear. Here, we performed integrated lipidomic and proteomic analyses of matched frontal (disease-vulnerable) and occipital (relatively spared) post-mortem cortices from individuals with genetic and sporadic FTLD-TDP, FTLD-tau (Pick’s disease, PiD), AD, and controls. FTLD and AD exhibited convergent lipid alterations, including reduced levels of cardiolipins and phosphatidylethanolamines, alongside increased gangliosides, diacylglycerols, cholesterol esters, acylcarnitines, and coenzyme Q, with generally greater changes in FTLD frontal cortex. FTLD displayed additional alterations, including reductions in bis(monoacylglycerol)phosphate, ceramides, phosphatidylserines, phosphatidylinositols, and sulfatides. These lipid changes were accompanied by proteomic alterations involving lysosomal proteins, phospholipases, phospholipid remodeling enzymes, and fatty acid oxidation pathways. Although lipidomic and proteomic signatures were broadly shared across FTLD subtypes, *GRN* associated FTLD-TDP and PiD showed the most extensive alterations. Triglycerides were selectively reduced in PiD in association with decreased DGAT1 expression, whereas cholesterol esters were elevated across all subtypes except *C9orf72* associated FTLD-TDP. These findings identify shared disruptions in lipid homeostasis and lysosomal lipid metabolism across FTLD and AD, highlighting convergent metabolic pathways underlying neurodegeneration.

## Introduction

Frontotemporal dementia (FTD) and Alzheimer’s disease (AD) are the most common forms of dementia before age 65 (1–3). Clinically, FTD and AD are distinct forms of neurodegenerative diseases. FTD exhibits regionally selective neurodegeneration prominently affecting the frontal, insular, and anterior temporal lobes (4–6). Most cases of FTD are associated with frontotemporal lobar degeneration (FTLD), characterized by either TDP-43 (FTLD-TDP) or tau (FTLD-tau) aggregation. FTLD-TDP and FTLD-tau are further classified into specific pathological subtypes, some of which are associated with both sporadic and familial disease. FTLD-TDP Type A (FTLD-TDP-A) occurs in sporadic cases and in individuals carrying with autosomal-dominant mutations in *GRN*, and *C9orf72* whereas FTLD-TDP Type C (FTLD-TDP-C) is primarily sporadic. In contrast, AD exhibits different anatomical vulnerability centered on the hippocampus and temporoparietal cortex (7). Despite clinical and anatomical differences, AD and FTLD share downstream consequences of neuroinflammation and neurodegeneration (4, 8).

Lipids are central to brain structure and function. The human cerebral cortex is particularly rich in sphingolipids, complex glycosphingolipids (e.g., gangliosides), and polyunsaturated phospholipids (e.g., phosphatidylethanolamine (PE) and plasmalogens) (9–11). Growing evidence implicates alterations in lipid pathways in FTLD. *GRN* haploinsufficiency, which is associated with FTLD-TDP-A, produces a characteristic lysosomal storage phenotype with ganglioside accumulation and loss of bis(monoacylglycerol)phosphate (BMP) (12–15), a phospholipid required for glycosphingolipid degradation. However, it is unclear whether similar lipid abnormalities occur in other forms of FTLD-TDP, including *C9orf72*-associated or sporadic FTLD-TDP-A, sporadic FTLD-TDP-C, or in FTLD-tau, such as Pick’s disease (PiD).

Similarly, AD is strongly linked to altered lipid metabolism. For instance, the *APOE4* allele, encoding a lipoprotein component, constitutes the strongest genetic risk factor for late-onset AD (16, 17) and exhibits altered cholesterol and sphingolipid metabolism, (18, 19).

Despite growing evidence that lipid dysregulation is a hallmark of both FTLD and AD, the molecular mechanisms underlying these alterations remain largely unknown. In particular, it is unclear whether lipid abnormalities are shared or accompanied by coordinated changes in proteins governing lipid metabolism, lysosomal function, and mitochondrial homeostasis. Furthermore, whether distinct FTLD subtypes converge on shared lipid-associated proteomic networks or exhibit subtype-specific molecular signatures is unresolved.

To address these questions, we performed mass spectrometry-based lipidomic and proteomic analyses of post-mortem human brain tissue from subjects with genetic (*GRN*- and *C9orf72*-associated) and sporadic FTLD-TDP-A, FTLD-TDP-C, Pick’s disease (PiD), a subtype of FTLD-Tau, AD, and neurologically normal controls. We analyzed both frontal cortex, a region prominently affected in FTLD, and occipital cortex, a comparatively spared region, enabling assessment of region-specific molecular remodeling.

This integrated multi-omic framework allowed direct comparison of shared and distinct molecular alterations across major neurodegenerative dementias and revealed convergent signatures of lysosomal and mitochondrial dysfunction, phospholipid depletion, sphingolipid remodeling, and altered lipid metabolism, alongside subtype-specific molecular changes associated with distinct genetic backgrounds.

## Results

### Study design

We analyzed post-mortem brain tissue samples using mass–spectrometry-based lipidomics and proteomics (**Fig. 1**). The tissue blocks were dissected from the middle frontal gyrus and lateral occipital cortex of six neurologically normal controls and patients with the following neuropathological diagnoses: 15 with FTLD-TDP-A*-GRN*, 4 with FTLD-TDP-A-*C9orf72*, 7 with sporadic FTLD-TDP-A, 15 with sporadic FTLD-TDP-C, 15 with FTLD-tau PiD, and 15 with AD. Clinical and neuropathological diagnoses were made using standard diagnostic criteria (20–23). Brains were donated with the consent of the participants or their surrogates in accordance with the Declaration of Helsinki, and the research was approved by the University of California, San Francisco Committee on Human Research. The demographic and clinical characteristics of the UCSF study cohort are summarized in the **Table 1 (Supplementary table 1)**.

**Fig. 1.**
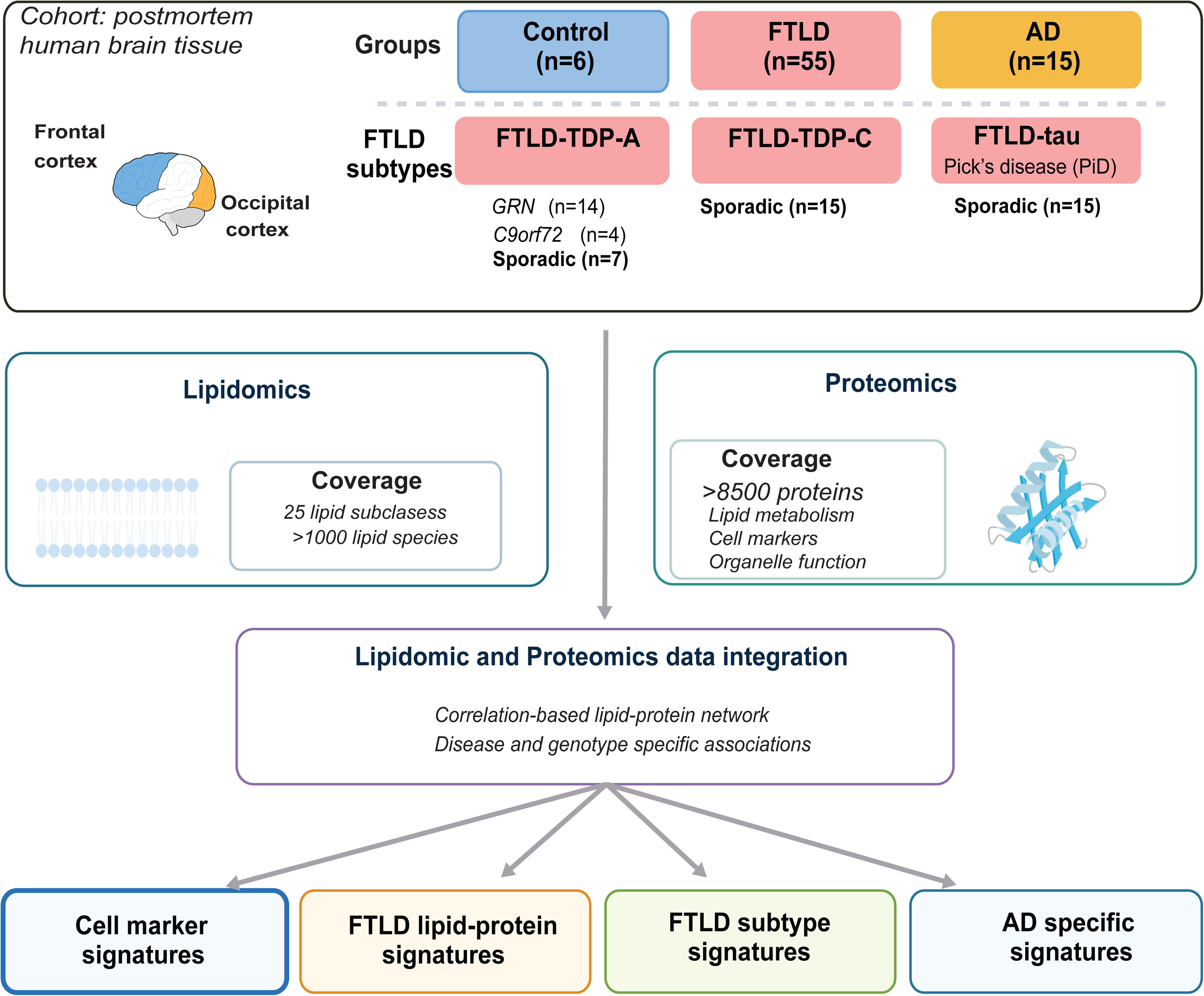
Study design for region- and disease-specific lipidomic and proteomic profiling of human brain. Postmortem frontal and occipital cortices from control, FTLD, and AD subjects were analyzed by LC-MS/MS-based lipidomics and proteomics. FTLD cases were stratified by genotype (*GRN, C9orf72*, and sporadic) and neuropathological diagnosis. Multi-omic integration enabled identification of region-, disease-, and subtype-specific lipidomic and proteomic alterations.

**Table 1.** Demographic and clinical characteristics of FTLD and AD cases stratified by genetic status and primary neuropathological diagnosis.

| Genetic Status | Primary Neuropathology | Clinical Diagnosis | N | Female (N) | Male (N) | Age at Death (yr, Mean ± SD) |
| --- | --- | --- | --- | --- | --- | --- |
| *GRN* | FTLT-DTP-A | bvFTD, nvPPA, CBS, mixed | 14 | 10 | 4 | 69.6 ± 8.1 |
| *C9orf72* | FTLT-DTP-A | bvFTD ± CBS | 4 | 1 | 3 | 67.3 ± 2.5 |
| Sporadic | FTLT-tau, Pick's disease | bvFTD, nvPPA, CBS, svPPA | 15 | 6 | 9 | 69.8 ± 5.6 |
| Sporadic | FTLT-DTP-A | bvFTD, CBS, PPA-mixed, AD vs FTD | 7 | 4 | 3 | 72.0 ± 6.6 |
| Sporadic | FTLT-DTP-C | svPPA | 14 | 9 | 5 | 69.9 ± 4.2 |
| Sporadic | AD | AD, lvPPA, PCA, AD vs svPPA, bvFTD (AD pathology) | 15 | 11 | 4 | 67.3 ± 6.0 |
| Controls | None | Control | 6 | 4 | 2 | 78.7 ± 11.2 |

Lipidomic analyses, with separate extraction methods for standard lipidomics and more polar glycolipids, led to the identification of more than 1,000 individual lipid species in more than 25 lipid subclasses **(Supplementary table 2)**. Proteomic coverage was deep and exceeded 8500 proteins per sample.

### Proteomics reveals cell-type composition changes across FTLD and AD subjects

The brain contains diverse cellular populations that can undergo substantial remodeling during neurodegeneration. To evaluate changes in cell composition of post-mortem brain tissue across FTLD and AD, we performed proteomic profiling and quantified neuron, microglia, astrocyte, and oligodendrocyte markers (**Fig. 2**). Neuronal markers were reduced across all FTLD subtypes, consistent with neurodegeneration or brain atrophy in this disease (**Fig. 2A**). In parallel, microglial and astrocytic markers were elevated in FTLD, indicating possibly glial hyper-activation and neuroinflammatory responses. These increases were observed across *GRN-, C9orf72-,* and sporadic FTLD-TDP and FTLD-tau groups and represented the most prominent proteomic changes.

**Fig. 2.**
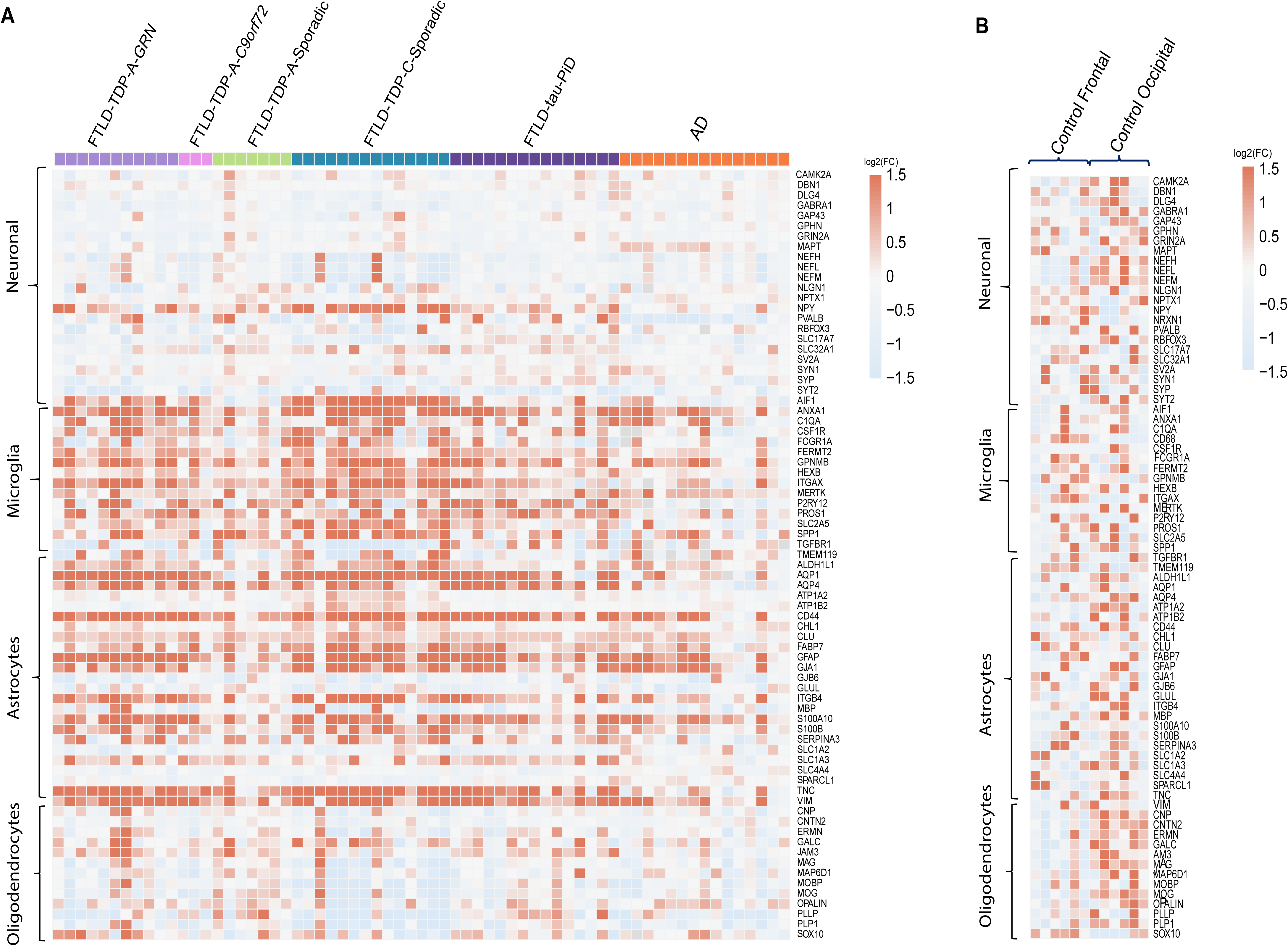
Proteomic analysis reveals cell-type-specific changes across FTLD and AD. **A** Heatmap of cell-type marker proteins in the frontal cortex across FTLD subtypes (*GRN, C9orf72,* and sporadic FTLD-TDP, and FTLD-tau) and AD, shown as log₂(FC) relative to controls. Markers are grouped by major brain cell types: neurons, microglia, astrocytes, and oligodendrocytes. **B** Heatmap of the cell-specific marker proteins in neurologically normal controls comparing frontal and occipital cortices, displayed as log₂(FC).

In AD, a similar pattern was characterized by reduced neuronal and elevated glial markers; however, the magnitude of these changes was comparatively weaker. Oligodendrocyte and myelination-associated markers generally tended to decrease in both FTLD and AD groups, although these reductions were more modest relative to neuronal and glial changes. These findings suggest that neuronal loss, accompanied by marked microglial and astrocytic activation, represents a major feature of FTLD pathology, while AD exhibits similar but less extensive proteomic remodeling.

The frontal and occipital control samples had comparable expression of neuronal and glial markers **(Fig. 2B)**, indicating that the observed changes in FTLD and AD reflect disease-specific cellular remodeling rather than regional differences in protein expression.

### Human frontal and occipital lobes of control subjects exhibit different lipid compositions

Frontal and occipital cortices of control subjects with no history of neurological or neurodegenerative disease were rich in glycerophospholipids and sphingolipids and had lower levels of neutral lipids. Specifically, ceramides (Cer), hexosylceramides (HexCer), PE, and plasmalogen (PE-O) were more abundant in the frontal cortex, whereas sphingomyelin (SM), glucosylsphingosine (GlcSph), sphingosine (SPH), sulfatides (ST), acylcarnitines (AcCa), diacylglycerols (DG), and lysophosphatidylserine (LPS) were comparatively reduced in frontal cortex in comparison to occipital cortex. Other lipid classes, such as PC, PS, CL or TG did not significantly differ between these brain regions (**Supplementary Fig. 1A& B**).

### FTLD brains exhibit lipid changes that are more marked in the frontal lobes

In previous studies of post-mortem FTLD brains, deficiency of the FTLD gene *GRN* resulted in low BMP levels and accumulation of gangliosides (12, 13), presumably indicating disrupted lysosomal sphingolipid catabolism. We tested whether these findings extend to other forms of FTLD by profiling lipids of frontal and occipital cortices from brains of individuals with different genetic or sporadic causes of FTLD and compared them with control samples.

Principal component analysis revealed separation between FTLD and control frontal cortex samples, indicating robust lipid changes in FTLD **(Fig. 3A**). Numerous lipid species were significantly changed, including gangliosides and neutral lipids that were increased, and phospholipids, such as BMP, that were decreased **(Fig. 3B, Supplementary Fig. 2A & B, and Supplementary table 3).**

**Fig. 3.**
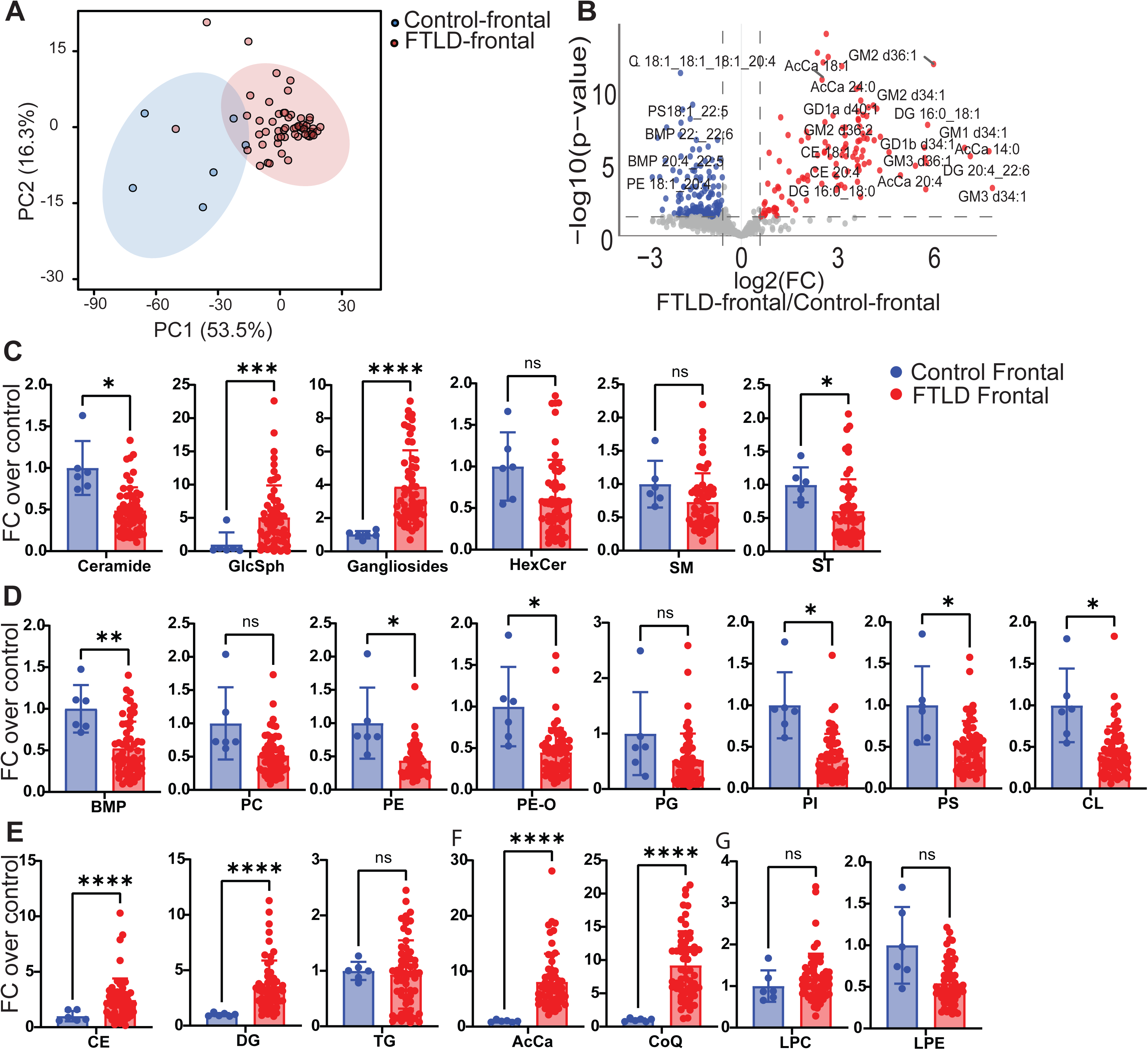
Lipid alterations in the frontal cortex of FTLD subjects. **A** Principal component analysis (PCA) of lipidome profiles from frontal cortex samples shows separation between control and FTLD groups; ellipses indicate 95% confidence intervals. **B** Volcano plot of differential lipid species in FTLD relative to controls, plotted as log₂(fold change) versus −log₁₀(p-value); significantly increased (red) and decreased (blue) species are highlighted (p < 0.05). **C–E** Lipid class–level comparisons in the frontal cortex. Bars represent mean fold-change (FC) relative to controls, and individual points represent lipid species within each class. Statistical significance was assessed between FTLD and control groups. P values were determined using an unpaired two-tailed Student’s t-test with Benjamini-Hochberg correction. *P < 0.05, **P < 0.01, ***P < 0.001, **** P<0.0001; ns, not significant.

FTLD frontal cortices exhibited marked alterations in sphingolipids (**Fig. 3C)**. GlcSph was significantly increased, and gangliosides were broadly elevated, consistent with disrupted ganglioside turnover. In contrast, ST and Cer were reduced.

Lipids of multiple glycerophospholipid classes were significantly reduced in FTLD frontal cortex **(Fig. 3D)**. These included PE, ether-linked PE (PE-O), PI, PS, and BMP, with the latter mirroring prior observations in *GRN*-associated FTLD (12). Decreases in cardiolipin (CL) and PE are consistent with previous studies suggesting mitochondrial stress (24).

In the neutral lipid classes, CE and DG were increased in FTLD frontal cortex, whereas TG levels were variable. In contrast, AcCa and coenzyme Q (CoQ) were elevated, while lysophospholipids exhibited more variability but were not different from controls **(Fig. 3F & G)**.

In the occipital cortex, lipid alteration changes were similar but milder (**Supplementary Fig. 3A & B**), indicating region-specific severity of lipid dysregulation, with the frontal cortex being a more affected region in FTLD, as expected. Numerous sphingolipids (e.g., Cer, ST, sphingosine, HexCer, and SM) were increased along with TGs. Glycerophospholipids did not change substantially in the occipital cortex.

### FTLD subtypes exhibit both shared and distinct lipidomic alterations

Comprehensive lipidomic profiling across the five FTLD subtypes, FTLD-TDP-A-*GRN*, FTLD-TDP-A-*C9orf72*, sporadic FTLD-TDP-A, sporadic FTLD-TDP-C and sporadic PiD identified both shared and subtype-specific changes **(Fig. 4A–F and supplementary Fig. 4A-D)**.

**Fig. 4.**
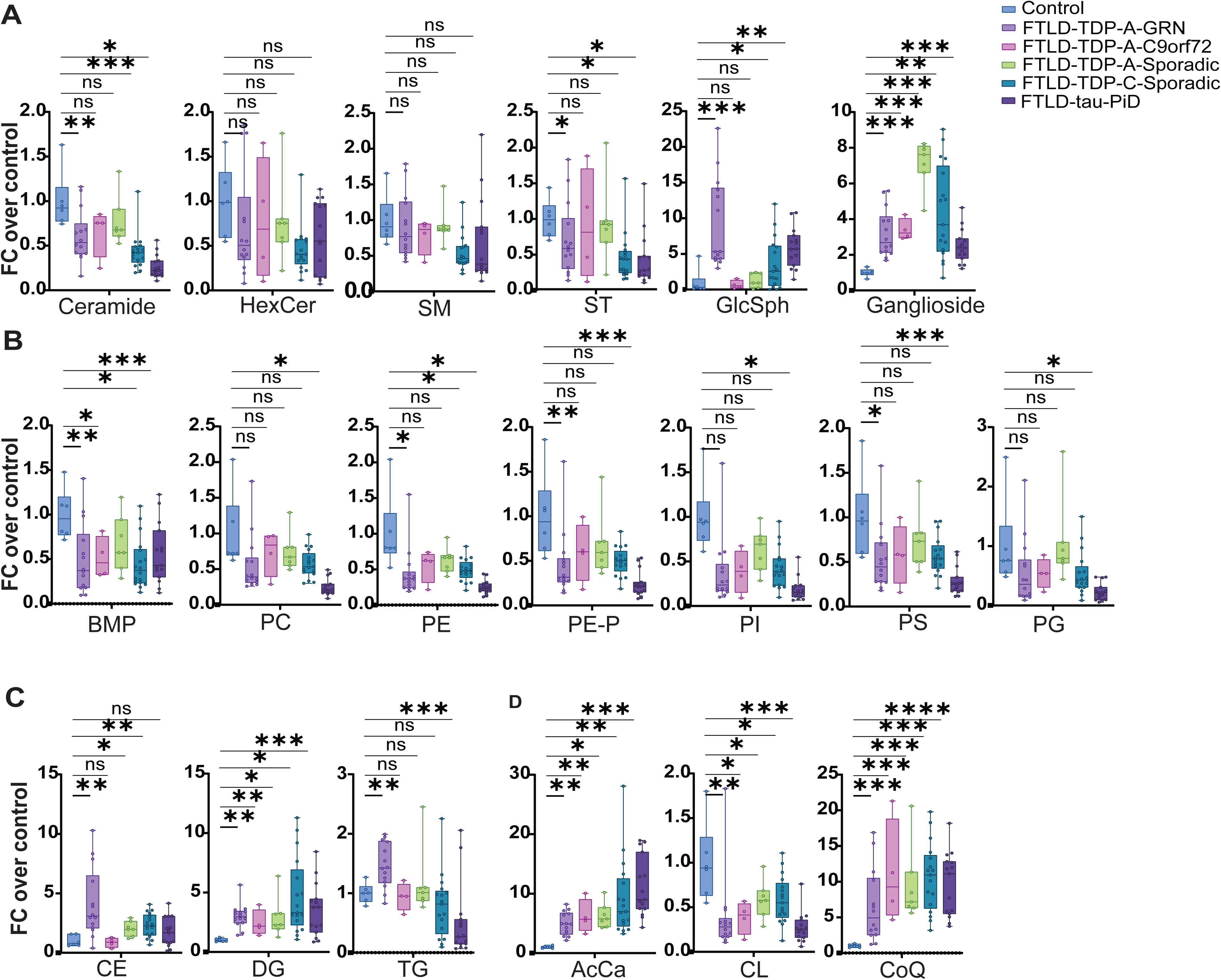
Genotype-specific lipid alterations in the frontal cortex in FTLD. Boxplots show fold-change of lipid levels relative to control across FTLD subtypes (*GRN, C9orf72,* and sporadic FTLD-TDP-A, FTLD-TDP-C, and FTLD- -tau PiD), with boxes indicating median and interquartile range and points representing individual lipid species. **A** Sphingolipid classes (Cer, HexCer, SM, ST, GlcSph, gangliosides). **B** Phospholipid classes (BMP, PC, PC-O, PE, PE-O, PI, PS, PG). **C** Neutral lipids and related classes (CE, DG, TG,). **D** AcCa, CL, CoQ. Each FTLD subtype was compared with controls using the Mann–Whitney U test, followed by Benjamini–Hochberg correction for multiple comparisons. *P < 0.05, **P < 0.01, ***P < 0.001, **** P<0.0001; ns, not significant.

Across all subtypes, the most consistent and prominent alteration was a marked depletion of BMP species, including BMP(18:0_18:1), BMP(18:1_20:4), and BMP(22:6_22:6). This conserved BMP reduction represents a central molecular signature of FTLD and implicates impaired lysosomal sphingolipid hydrolysis in disease etiology. This loss was accompanied by reductions in mitochondrial lipids, particularly cardiolipins, and decreases in PI, PE, PE-O, and PS, supporting a broader deficit in membrane phospholipid homeostasis. DG and AcCa were elevated across subtypes, suggesting impaired fatty acid oxidation and mitochondrial stress **(Fig. 4B–F and Supplementary Fig. 4A & F)**. In parallel, all FTLD subtypes exhibited accumulation of gangliosides, possibly due to reduced BMP levels **(Fig. 4A & B)**.

Despite the shared abnormalities, each FTLD subtype also exhibited distinct lipid changes. FTLD-TDP-A-*GRN* had the most pronounced and widespread lipid perturbations. Particularly, GlcSph was increased much more than all other groups, and gangliosides were markedly elevated. CE and TG demonstrated a strong increase. Other sphingolipid classes such as Cer, HexCer, and SM were unchanged (**Fig 4A**).

FTLD-TDP-A-*C9orf72* was represented by a small cohort (n=4). Despite its small sample size, this cohort had a consistent trend of a milder lipid phenotype than the other subtypes. While BMP and cardiolipins were reduced, several other glycerophospholipid classes that were decreased in other subtypes showed less apparent reductions at the class level in samples of *C9orf72* subjects. Most other lipid classes remained close to control levels, with only increases in gangliosides, DG, and selected AcCa (**Fig A, C &D**).

FTLD-tau PiD, demonstrated a distinctive reduction in sphingolipids. Cer, HexCer, SM, and ST were lower than other FTLD subtypes. CE elevation was limited, and TG levels appeared significantly reduced.

The two sporadic FTLD-TDP subtypes shared a similar lipid pattern, marked by robust elevations in CE and AcCa, particularly in FTLD-C.

### Regional and disease-subtype specific proteomic alterations involved in lipid metabolism in FTLD

Principal component analysis of frontal cortex proteomes showed separation between FTLD and controls, indicating distinct proteomic profiles **(Supplementary Fig. 5A and Supplementary table 4)**. Differential expression analysis revealed upregulation of proteins involved in lipid metabolism and lysosomal function, including NPC2, PLD2, ASAH1 and LIPA, alongside with enzymes linked to fatty acid metabolism and oxidative stress such as ACADL. In contrast, proteins involved in phospholipid synthesis and membrane maintenance, including LPCAT4 and PHOSPHO1, were reduced **(Fig. 5A)**. Pathway enrichment analysis supported these findings, highlighting upregulation of lipid-related and degradative processes, including PPAR signaling, cholesterol metabolism, lysosome, peroxisome, and fatty acid metabolism pathways **(Fig. 5B).** In contrast, downregulated proteins were enriched in ether lipid metabolism and glycerophospholipid metabolism **(Fig. 5C)**, indicating impaired membrane lipid synthesis and remodeling.

**Fig. 5.**
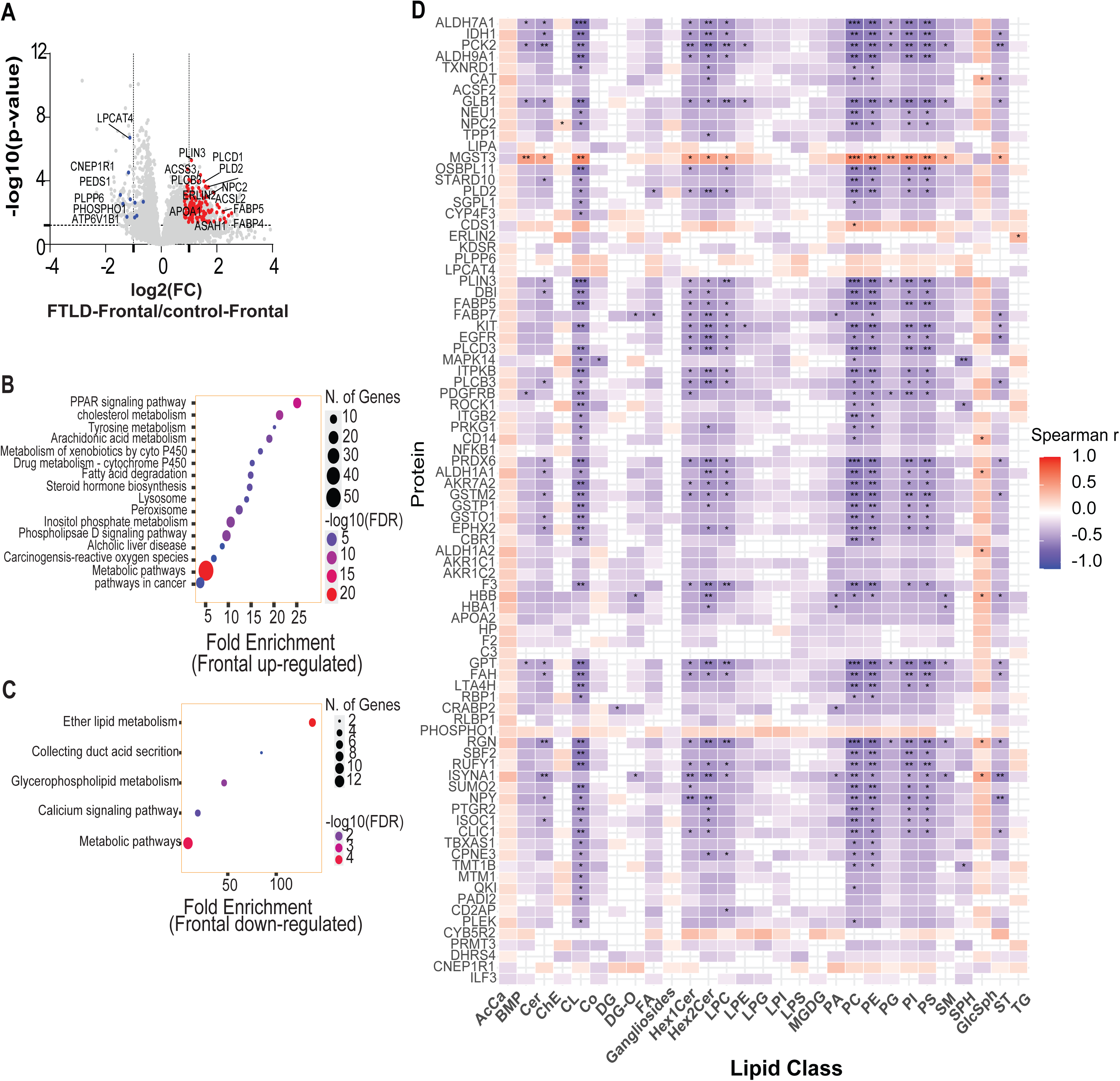
Integrated proteomic and lipidomic remodeling of the frontal cortex in FTLD. **A** Volcano plot of differential protein abundance in the frontal cortex comparing FTLD to controls, displayed as log₂(fold-change) versus −log₁₀(*p*-value). Significantly upregulated (red) and downregulated (blue) proteins (*p*<0.05) are highlighted, with selected proteins annotated. **B** Pathway enrichment analysis of proteins upregulated in the frontal-cortex in FTLD, with enriched biological pathways plotted by fold enrichment and −log₁₀(FDR). **C** Pathway enrichment analysis of proteins downregulated in the frontal cortex in FTLD, highlighting pathways related to lipid metabolism, membrane organization, and cellular signaling. **D** Heatmap of Spearman correlations between lipid classes and lipid metabolism-related proteins in FTLD frontal cortex samples. Significant associations are indicated by *, **, and *** P < 0.001 (|Spearman r| ≥ 0.3, p < 0.05).

To further investigate relationships between altered lipid classes and lipid metabolism-associated proteins within disease states, we performed Spearman correlation analysis exclusively in FTLD frontal cortex samples using a curated set of proteins involved in lipid metabolism, lysosomal pathways, membrane remodeling, oxidative stress, and mitochondrial function (**Fig. 5D**). Lysosomal proteins, including NPC2, GLB1, NEU1, TPP1, and LIPA, showed similar correlation tendencies across several sphingolipid and phospholipid classes, whereas ER- and membrane-associated proteins such as ERLIN2, SGPL1, KDSR, LPCAT4, STARD10, PLD2, and MGST3 displayed related lipid association patterns. Mitochondrial and metabolic proteins, including ALDH7A1, IDH1, PCK2, ACSF2, TXNRD1, and CAT, generally exhibited negative correlations with multiple phospholipid classes. Lipid droplet-associated proteins, including PLIN3, FABP5, FABP7, and DBI, also demonstrated related lipid association profiles. Stronger significant correlations (|Spearman r| ≥ 0.3, p < 0.05) were primarily observed within sphingolipid, glycerophospholipid, and ether lipid classes, supporting widespread disruption of metabolic homeostasis in FTLD frontal cortices.

FTLD subtypes had both shared and distinct proteome alterations **(Fig. 6A–E).** *GRN-, MAPT-*, and sporadic FTLD showed widespread dysregulation of lipid-associated proteins, whereas changes were less pronounced in *C9orf72*-associated diseases, with a general pattern of increased lipid metabolism and lysosomal proteins alongside reduced phospholipid-associated enzymes.

**Fig. 6.**
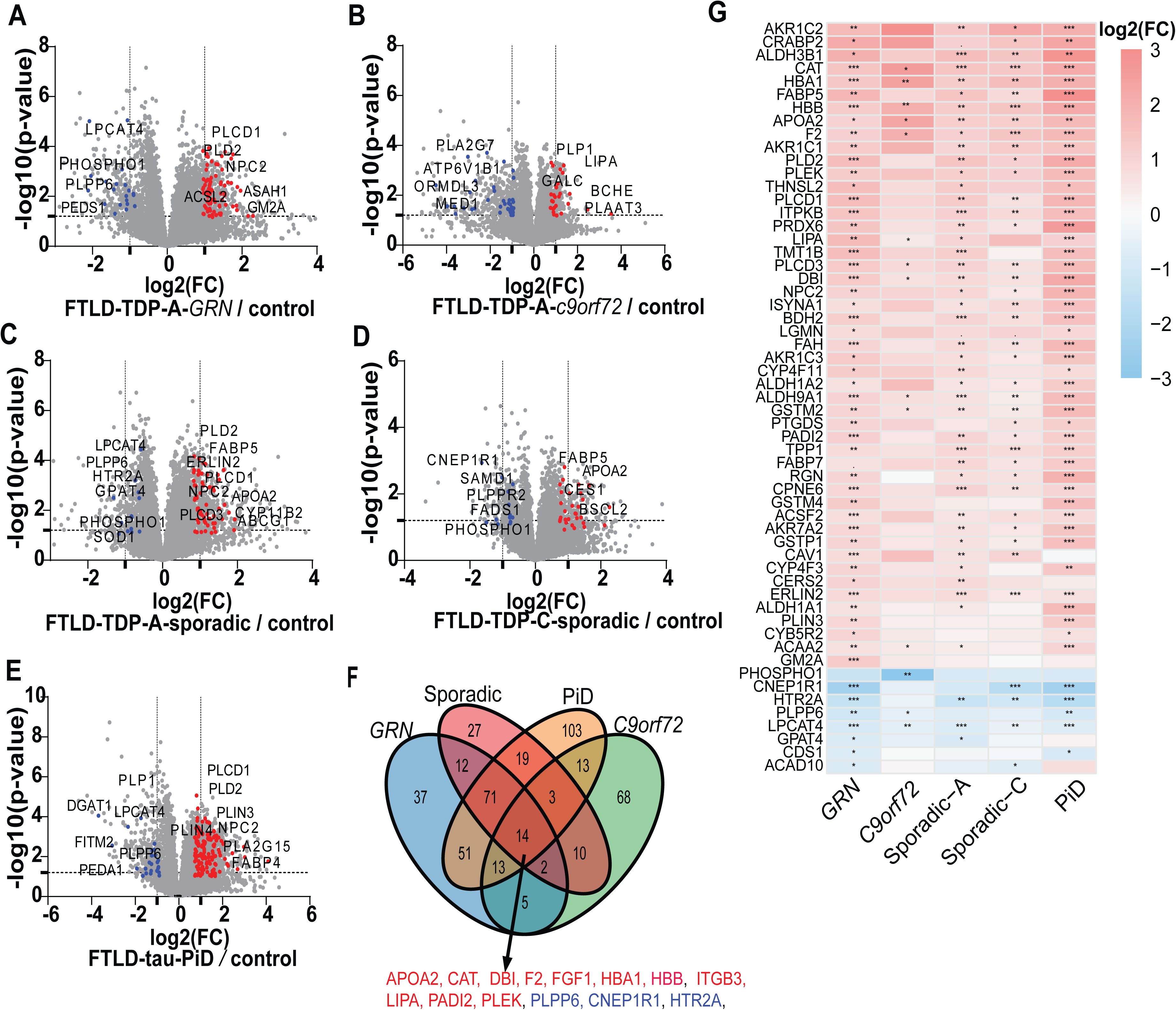
Shared and FTLD subtype-specific proteomic alterations. **A–E** Volcano plots showing differential protein expression in frontal cortex for FTLD subtypes (FTLD-TDP-A-*GRN,* FTLD-TDP-A-*C9orf72,* sporadic FTLD-TDP-A, FTLD-TDP-C, and FTLD-tau PiD) relative to controls. Significantly upregulated proteins are shown in red and downregulated proteins in blue. Selected proteins are labeled. **F** Venn diagram illustrating overlap of differentially expressed proteins across FTLD subtypes, highlighting shared and subtype-specific signatures. **G** Heatmap of log₂(fold-change) for selected proteins across FTLD subtypes. Color scale indicates magnitude of change, and asterisks denote statistical significance.

A core set of 14 proteins was consistently altered across subtypes **(Fig. 6F; Supplementary Table 3),** defining a signature of lipid remodeling and cellular stress. This included regulators of lipid turnover (LIPA, PLPP6, CNEP1R1, DBI) and proteins liked to oxidative stress and inflammation (CAT, APOA2, F2).

These patterns are reflected in the heatmap **(Fig. 6G)**, which shows coordinated upregulation of lipid metabolism and lysosomal proteins and downregulation of phospholipid-related proteins across subtypes. The magnitude of change was greatest in *GRN* and sporadic FTLD-TDP, and FTLD-tau PiD, and attenuated in FTLD-TDP-A-*C9orf72*.

Subtype-specific differences were also evident: *GRN*-FTLD showed stronger lysosomal enrichment (e.g., GM2A), FTLD-tau PiD displayed broader alterations including fatty acid and mitochondrial metabolism (e.g., ACADL, DGAT1), and sporadic FTLD showed changes in cholesterol transport and oxidative pathways (e.g., ABCA7, EPHX1).

Finally, we extended our analysis to the occipital cortex. PCA showed separation between FTLD and control samples **(Supplementary Fig. 5B)**, but differential expression analysis identified fewer lipid-related proteins, including APOE and APOC1, along with inflammatory markers **(Supplementary Fig. 5C)**. Pathway enrichment highlighted alterations in cholesterol and lipid metabolism **(Supplementary Fig. 5D)**, indicating less extensive lipid-associated proteomic changes than in the frontal cortex.

### AD exhibits both shared and distinct lipidomic and lipid-associated proteomic alterations

We examined whether lipid alterations observed in FTLD were also present in AD. In AD frontal cortices, lipid remodeling was present but comparatively selective **(Fig. 7A–C; Supplementary Fig. 6A–B)**. Among sphingolipids, ceramide, hexosylceramides, sphingomyelin, and sulfatides were largely unchanged, whereas gangliosides were increased, driven primarily by GM1 and GM3 species **(Fig. 7A, Supplementary Fig. 6B).**

**Fig. 7.**
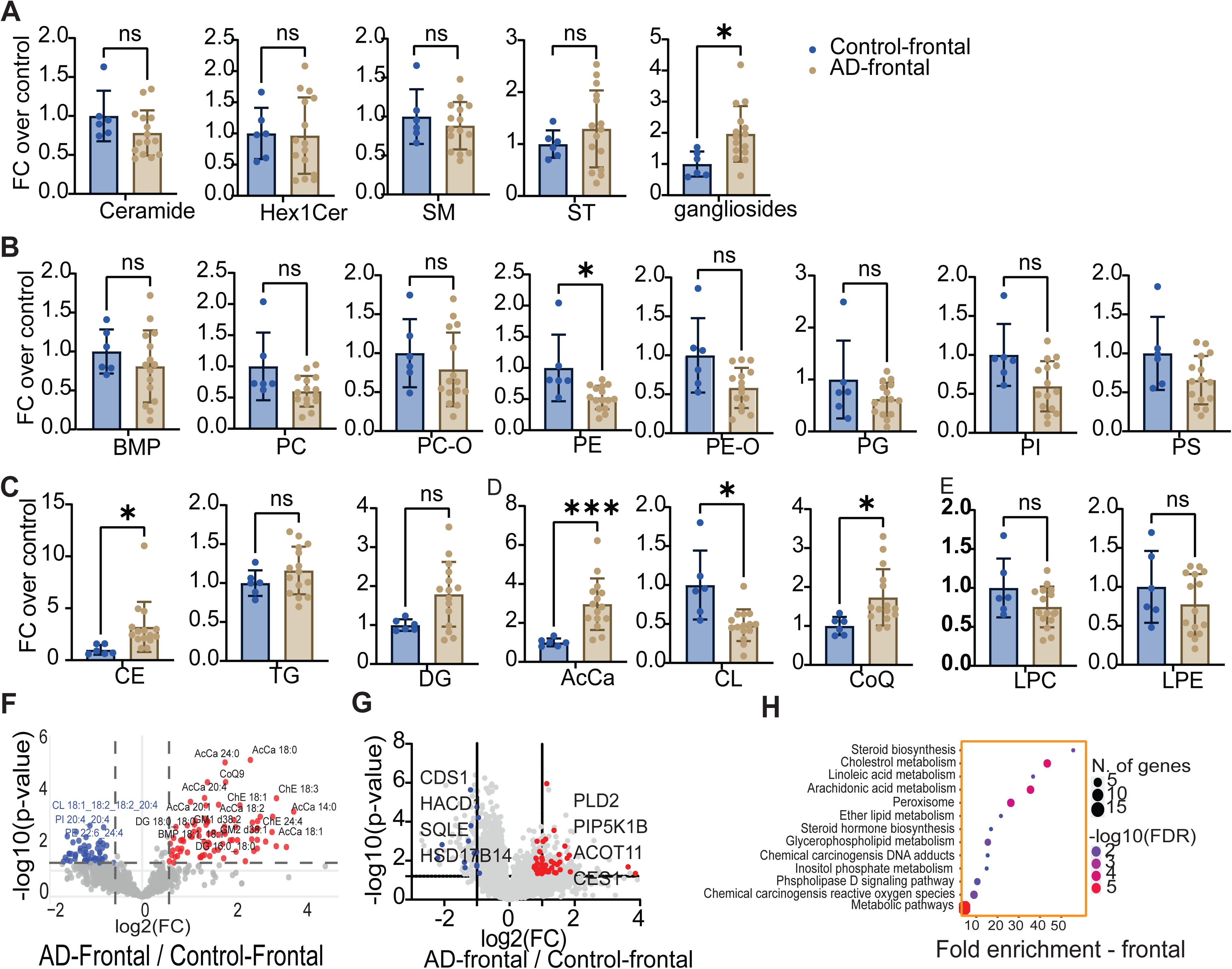
Lipid and protein alterations in the frontal cortex of AD subjects. **>A–E** Lipid class–level comparisons between control and AD frontal cortices across sphingolipids, phospholipids, and neutral/mitochondrial-related lipids. Boxplots show fold change relative to control frontal cortex, with bars indicating mean ± SD and points representing individual lipid species. P values were determined using an unpaired two-tailed Student’s t-test with Benjamini–Hochberg correction. *P < 0.05, ***P < 0.001; ns, not significant. **F** Volcano plot of differential lipid species in the frontal cortex comparing AD to controls, log₂(fold-change) vs −log₁₀(p-value), with significantly increased species highlighted in red and significantly decreased species highlighted in blue. **G** Volcano plot of differential protein abundance in the frontal cortex comparing AD to controls, log₂(fold-change) vs −log₁₀(p-value), with significantly increased species highlighted in red and significantly decreased species highlighted in blue. **H** Pathway enrichment analysis of lipid-related, upregulated proteins in the frontal cortex in AD, with enriched biological pathways plotted by fold enrichment and −log₁₀(FDR).

Glycerophospholipid changes were limited, with reductions observed mainly in cardiolipin and PE, while other classes, including BMP, PI, PS, and PE-O, trended lower but were not significantly altered **(Fig. 7B)**. Neutral lipids showed selective increases, with CE most prominently elevated, and DG showed modest changes and TG remained largely unchanged (**Fig. 6C**). AcCa and CoQ were increased **(Fig. 7D).**

At the protein level, these changes were accompanied by reduced abundance of enzymes involved in phospholipid synthesis and elongation (e.g., CDS1, HACD2, FA2H, SQLE) and increased levels of proteins associated with lipid turnover, transport, and fatty acid metabolism (e.g., PLD2, CES1, STARD3, PLTP, CYP4F2, AKR1C2). Pathway enrichment analysis highlighted cholesterol and glycerophospholipid metabolic pathways **(Fig. 7C–H).**

Compared with FTLD, lipid alterations in AD were less extensive, with fewer affected glycerophospholipids and more restricted sphingolipid changes. Consistent with this, fewer lipid-associated proteins were differentially expressed in AD than in FTLD. Despite these quantitative differences, both diseases converged on a shared core of perturbed proteins involved in pglycerohospholipid metabolism and lipid signaling (CRABP2, AKR1C2, PLD2, CDS1, EPHX2), defining a conserved lipid-linked cellular stress framework common to neurodegeneration

## Discussion

In this study, we integrated lipidomic and proteomic profiling of post-mortem brain samples to define region- and subtype-specific metabolic alterations across FTLD and AD. Several major conclusions emerge. First, FTLD is characterized by consistent lipid dysregulation across genetic and sporadic forms, most prominently in the frontal cortex for FTLD, including reductions in many membrane glycerphospholipids together with increases in gangliosides, glucosylsphingosine, neutral lipids, and lipids associated with mitochondrial dysfunction. Second, these lipid changes are accompanied by proteome changes involving glycerophospholipid metabolism and lysosomal pathways, including reduced levels of glycerophospholipid remodeling enzymes (e.g., LPCAT4) with increased phospholipases (e.g., PLD2), alongside increased levels of lysosomal lipid-handling proteins (e.g., NPC2, LIPA). Third, AD brains share features of this lipid dysregulation pattern, including reduced glycerophospholipids, and increased gangliosides and neutral lipids, although these alterations were generally less pronounced than in FTLD. This difference may reflect the use of frontal cortex samples rather than more severely affected temporoparietal regions typically involved in AD.

Across FTLD subtypes, we observed a conserved lipid signature characterized by reduced levels of glycerophospholipids (including CL, BMP, PE, PE-ethers, PI, PS), together with accumulation of gangliosides, DG, AcCa, and CoQ. Reductions in glycerophospholipids may partly reflect underlying neurodegeneration, including loss of neurons and oligodendrocytes, as supported by the decreased cellular markers observed in this study and in prior reports (25–28).

Lysosomal dysfunction may be a shared disease mechanism across different forms of FTLD. The consistent decrease in BMP, a lipid enriched in late endosomes and lysosomes, together with increased glycosphingolipids suggests impaired lysosomal lipid degradation across FTLD subtypes (13, 25, 27, 29, 30). This notion is supported by proteomic changes showing increased abundance of lysosomal and lipid turnover proteins (e.g., NPC2, ASAH1, TPP1) and phospholipases (e.g., PLD2, PLCD1).

In parallel, mitochondrial alterations appear to be a second shared feature across FTLD subtypes. Reduced cardiolipin and increased AcCa and CoQ suggest impaired mitochondrial function and impaired fatty acid oxidation (31, 32), consistent with changes in mitochondrial enzymes (e.g., ACADL). Together, these findings support a model in which lysosomal lipid accumulation, mitochondrial stress, and reduced glycerophospholipids, possibly reflecting cell loss, occur in parallel in FTLD.

While shared features were evident, subtype-specific differences highlight distinct disease mechanisms. *GRN*-FTLD-TDP-A showed pronounced lysosomal changes, including increased GM2A, a lysosomal lipid activator involved in glycosphingolipid degradation. These changes suggest a compensatory upregulation in response to ganglioside accumulation (33–36). FTLD-tau PiD exhibited broader metabolic alterations, including reduced triglycerides associated with decreased levels of DGAT1 (synthesizing TG) and changes in fatty acid metabolism (e.g., ACADL), enzymes involved in mitochondrial β-oxidation, consistent with increased lipid utilization and metabolic demand. (37–39). Sporadic FTLD brains had altered proteins involved in cholesterol transport and oxidative pathways (e.g., ABCA7, GSTM4) (40). In contrast, *C9orf72*-FTLD-TDP-A showed attenuated changes with no significant alteration for instance in NPC2, suggesting minimal disruption of lysosomal cholesterol metabolism (11).

AD exhibits partial overlaps with this lipid program, including accumulation of CE, gangliosides, AcCa, and CoQ, as well as increased levels of proteins involved in cholesterol transport. These findings are consistent with prior work linking *APOE*-dependent lipid trafficking and sterol metabolism to AD pathogenesis (26, 41). At the protein level, these changes are accompanied by alterations in lipid transport and turnover proteins (e.g., PLD2, CES1), consistent with modified lipid handling. However, in contrast to FTLD, AD samples exhibited no significant alteration in BMP levels and did not show the same phospholipid reduction or coordinated lysosomal–membrane disruption. These findings suggest that while lysosomal and lipid trafficking pathways are perturbed in both diseases, the degree and nature of disruption might differ. Importantly, these observations are based on samples from the frontal cortex, which is relatively less affected in early AD than the temporoparietal regions. Therefore, the absence of more pronounced phospholipid alterations in AD should be interpreted within this regional context (42). A further limitation of the current study is that only few samples are available for some of the subtypes. While this highlights the robustness of the most significant lipid and protein changes, it suggests further changes may be uncovered as more samples are analyzed.

In summary, our findings show that altered lipid metabolism is a prominent feature of both FTLD and AD brains. Across these diseases, brain tissue shows reduced glycerophospholipids, consistent with loss of neurons and oligodendrocytes, and accumulation of gangliosides, suggesting endo-lysosomal dysfunction, and reduced cardiolipin, consistent with mitochondrial impairment. Proteomics for both diseases also feature changes in lipid metabolism proteins. These changes support the hypothesis that lipid alterations are closely associated with neurodegeneration and may contribute to disease pathogenesis.

## Methods

### Human brain samples

Human brain tissue samples were obtained from the Neurodegenerative Disease Brain Bank at the University of California, San Francisco. In keeping with the guidelines put forth in the Declaration of Helsinki, patients or their surrogates provided informed consent for brain donation prior to autopsy, Neuropathological diagnoses were made following consensus diagnostic criteria (20, 43) using previously described histological and immunohistochemical methods (Kim et al., 2012; Tartaglia et el., 2010). Genetic screening was performed for common and rare FTLD-related genes as previously described (43). Cases were identified based on neuropathological diagnosis and genetic screening. Fresh-frozen blocks of the middle frontal gyrus and lateral occipital cortex were cut from coronal slabs and stored at -80C for further analyses.

### Chemicals

Liquid chromatography–mass spectroscopy (LC–MS)-grade chloroform (MilliporeSigma, Cat# 650498), methanol (Fisher Chemical, Cat# A452), isopropanol (IPA) (Fisher Chemical, Cat# A461), acetonitrile (ACN) (Fisher Chemical, Cat# A955), water (Fisher Chemical, Cat# W6), formic acid (Thermo Fisher Scientific, Cat# 85178), acetic acid (MilliporeSigma, Cat# 695092), ammonium formate (MilliporeSigma, Cat# 70221), ammonium acetate (MilliporeSigma, Cat# A1542), trifluoroacetic acid (Thermo Fisher Scientific, Cat# 28904), and ammonium bicarbonate (MilliporeSigma, Cat# 09830) were used for lipidomic and proteomic analyses. SPLASH LIPIDOMIX internal standards (Avanti Polar Lipids, Cat# 330707) and GM3-d5 standards (Matreya LLC) were used for normalization. SOLA HRP solid-phase extraction cartridges (Thermo Scientific, Cat# 60109-001) and Sep-Pak cartridges (Waters) were used for sample cleanup. Protein digestion reagents included urea (MilliporeSigma, Cat# U5128), tris (2-carboxyethyl)phosphine) (Thermo Fisher Scientific, Cat# 77720), iodoacetamide (MilliporeSigma, Cat# I1149), dithiothreitol (Bio-Rad, Cat# 1610611), Lys-C protease (Wako Chemicals, Cat# 129-02541), and sequencing-grade trypsin (Promega, Cat# V5111).

### Lipid extraction

For lipidomic analysis, weighed brain tissue was homogenized in ice-cold phosphate-buffered saline with a bead mill homogenizer. Tissue lysates (∼50 μg protein) were transferred into Pyrex glass tubes with PTFE-lined caps. Lipids were extracted using a modified Folch method (44). Briefly, 6 mL of ice-cold chloroform: methanol (2:1, v/v) and 1.5 mL of water were added to each sample and vortexed thoroughly to ensure complete phase mixing. SPLASH LIPIDOMIX internal standards (Avanti Polar Lipids) were added prior to extraction for normalization. Samples were centrifuged at 1,000 rpm for 20 min at 4°C to achieve phase separation. The lower organic phase was carefully collected using glass pipettes, avoiding the interphase containing proteins and debris. Extracted lipids were dried under a nitrogen stream and reconstituted in 100 μL of IPA:ACN:water (60:35:5, v/v/v). Samples were stored at −80°C until LC–MS/MS analysis.

### Ganglioside extraction

For ganglioside analysis, ∼100 μg of homogenized tissue was extracted with methanol and centrifuged at 1,000 × g for 20 min to pellet proteins. The supernatant containing gangliosides was transferred to fresh glass tubes and dried under nitrogen. Dried extracts were reconstituted in 1 mL of LC-MS grade water and desalted using SOLA HRP solid-phase extraction cartridges (Thermo Scientific, 30 mg/2 mL). Cartridges were pre-conditioned with methanol (3 × 1 mL) and equilibrated with water (3 × 1 mL). Samples were loaded, washed with 2 mL of water, and gangliosides were eluted with 3 mL of methanol. Eluates were dried under nitrogen and reconstituted in methanol:water:chloroform (60:9:120, v/v/v) for LC–MS/MS analysis.

### LC-MS/MS analysis of lipids

Lipid extracts were analyzed using ultra-high-performance liquid chromatography (UHPLC) coupled to tandem mass spectrometry (MS/MS). Separation was performed on a C30 reverse-phase column (Thermo Acclaim C30, 2.1 × 150 mm, 2.6 μm) maintained at 50°C using a Vanquish Horizon UHPLC system (Thermo Fisher Scientific) coupled to an Orbitrap Exploris 240 MS equipped with a heated electrospray ionization source.

Samples (2 μL) were analyzed in both positive and negative ionization modes. Mobile phase A consisted of water:ACN (60:40, v/v) with 10 mM ammonium formate and 0.1% formic acid. Mobile phase B consisted of IPA:ACN (90:10, v/v) with identical additives.

The gradient was as follows: 30% B (−3 to 0 min), increased to 43% B (0–2 min), 55% B (2–2.1 min), 65% B (2.1–12 min), 85% B (12–18 min), and 100% B (18–20 min), held at 100% B (20–25 min), and followed by re-equilibration to 30% B (25.1–28 min). The flow rate was 0.26 mL/min.

MS parameters included an ion transfer tube temperature of 300°C and vaporizer temperature of 275°C. Spray voltages were set to 3.25 kV (positive) and 2.5 kV (negative). MS1 resolution was 120,000 and MS2 resolution was 30,000. Data-dependent acquisition was performed with a cycle time of 1.5 s, isolation window of 1 m/z, and dynamic exclusion of 2.5 s. HCD fragmentation energies were stepped at 15%, 25%, and 35%. Full-scan acquisition covered an m/z range of 250–1700. Internal calibration was performed using EASY-IC™.

### LC-MS/MS analysis of gangliosides

Gangliosides were analyzed using a Vanquish UHPLC system coupled to an Orbitrap Exploris 240 MS. Separation was achieved on a Kinetex HILIC column (Phenomenex, 2.6 μm, 100 × 2.1 mm) at 50°C. Mobile phase A consisted of ACN with 0.2% acetic acid, and mobile phase B consisted of water containing 10 mM ammonium acetate (pH 6.1). A linear gradient was applied from 12.3% B to 22.1% B over 15 minutes, followed by column re-equilibration. The flow rate was 0.6 mL/min. MS was performed in negative ion mode with a spray voltage of −4.5 kV, capillary temperature of 300°C, and vaporizer temperature of 250°C. MS1 resolution was 120,000 (m/z range 700–1800), and MS2 resolution was 30,000. Internal calibration was performed using EASY-IC™.

### Data processing and quantification

Raw LC–MS/MS data were processed using LipidSearch 5.1 with precursor and product ion tolerances of 5 ppm and 8 ppm, respectively. Lipid identifications were aligned across samples and filtered based on signal quality and reproducibility, followed by additional normalization and aggregation using an in-house pipeline (Lipidcruncher (45)). Quantification was performed by normalizing lipid peak areas to class-specific internal standards and total protein content. Gangliosides were quantified relative to GM3-d5 standards.

Quality control was ensured using internal standards to monitor extraction efficiency, signal consistency, and instrument performance, including retention time stability and mass accuracy. Reproducibility was assessed by retaining lipid species consistently detected across samples and evaluating variation across replicates. Principal component analysis and clustering were used to confirm sample grouping and identify potential outliers or batch effects.

Lipid identifications were validated based on accurate mass, MS/MS fragmentation, and database matching, and only high-confidence species were included in downstream analyses. Consistency between lipidomic and proteomic changes was used as an additional layer of validation, supporting the robustness of the observed biological patterns.

### Proteomics analyses

#### Protein digestion

Samples were diluted 1:1 with 8M urea, 50 mM EPPS (pH 8.5). Following protein quantification using the Pierce bicinchoninic acid (BCA) assay (Thermo Fisher), 100 µg of proteins was reduced with tris 2-(carboxyethyl) phosphine to a final concentration of 5 mM for 20 min at room temperature (RT) with shaking at 1,000 rpm on a thermomixer. Free cysteine residues were then alkylated with iodoacetamide at a final concentration of 10 mM for 30 minutes at RT in the dark. Alkylation was quenched with dithiothreitol (DTT) to a final concentration of 5 mM and incubated for 15 minutes at RT. Lys-C was added at a 1:200 enzyme-to-protein ratio, and the samples were incubated for 2 hours at 25°C with shaking at 1,000 rpm. Samples were then diluted with 50 mM ammonium bicarbonate to reduce the urea concentration to 2M, and trypsin was added at a 1:100 enzyme-to-protein ratio. The samples were incubated overnight at 37°C with shaking at 1,150 rpm.

After digestion, the samples were acidified to pH <3 by adding 50% trifluoroacetic acid (TFA) and desalted using Sep-Pak cartridges (Waters) conditioned with 100% MeOH, 70% ACN/0.1%TFA, 5% ACN/0.1% TFA twice. After sample loading, the cartridges were washed with 5% ACN/0.1% formic acid (FA) twice and the peptides eluted with 70% ACN/0.1% FA twice. Desalted peptides were dried under vacuum in a SpeedVac and reconstituted in 12 µL of 0.1% FA.

#### MS analyses

Peptides were separated using a trap-and-elute method on a 25 cm, 75 µm column packed with 1.7 µm C18 particles (Aurora Ultimate IonOpticks). Peptide separation was performed over a 54.2-minute gradient as follow: 3-5% mobile phase B over 25 seconds; 5-6% B over 30 seconds, 6–7% B over 25 seconds, 7–22% B over 34 min, 22–36 %B over 11 min, 36–95% B over 1 min, followed by a 6-min hold at 95% B and re-equilibration to 3% B over 1 min at a flow rate of 300 nL/min. The mobile phase A consisted of 0.1% FA in HPLC-grade water, and mobile phase B consisted of 90% ACN with 0.1% FA. Separation was performed using an EASY-nLC 1200 liquid chromatography system (Thermo Fisher Scientific).

MS data were acquired on an Orbitrap Astral mass spectrometer (Thermo Fisher Scientific) in a data-independent acquisition (DIA) mode with a normalized collision energy of 25%. MS1 spectra were acquired in the Orbitrap at a resolution of 240 K, with a normalized AGC target of 500%, a custom maximum injection time, and a scan range of 380–980 m/z. MS/MS spectra were acquired in the Astral analyzer using a 2 m/z isolation window, a scan range of 150–2000 m/z, a precursor mass range of 380–980 m/z, and a loop cycle time of 0.6 seconds.

#### <u>D</u>ata <u>I</u>ndependent <u>A</u>cquisition (DIA) mass spectrometry data analysis

The raw data files were processed using Spectronaut version 19.6 (Biognosys) and searched with the PULSAR search engine against a *Homo Sapiens* UniProt protein database downloaded on 2024/08/09 (42,524 entries). Cysteine carbamidomethylation was specified as a fixed modification, while methionine oxidation, acetylation of the protein N-terminus, and deamidation (NQ) were set as variable modifications. A maximum of two trypsin missed cleavages was permitted. The searches utilized a reversed sequence decoy strategy to control the peptide false discovery rate (FDR), with a threshold of 1% FDR set for identification. An unpaired t-test was used to calculate the p-value in the differential analysis, and a volcano plot was generated based on log2 fold change (log2FC) and q-value (multiple testing corrected p-value using the Benjamini-Hochberg method). A q-value of ≤0.05 was considered the statistically significant cut-off.

## Data availability

The mass spectrometry proteomics raw data generated in this study have been deposited in the ProteomeXchange Consortium via the PRIDE partner repository under accession code identifier PXD080575 (46). Lipidomics raw data have been deposited in Metabolomics Workbench under identifier ST004317 (47).

## Acknowledgments

We thank members of Farese & Walther for helpful discussions and Gary Howard for editorial assistance. This work was supported by a grant from the Bluefield Project to Cure FTD (to R.V.F. and T.C.W.), and postdoctoral fellowship grants from the Bluefield Project to Cure FTD (to Y.A.). T.C.W. is a Howard Hughes Medical Institute Investigator. We acknowledge NIH/NCI Cancer Center Support Grant (Core grant P30 CA008748) to MSKCC and the Lipidomics Innovation laboratory at MSKCC. The UCSF Neurodegenerative Disease Brain Bank receives funding support from NIH grants P30AG062422, P01AG019724, U01AG057195, and U19AG063911, as well as the Rainwater Charitable Foundation and the Bluefield Project to Cure FTD. The authors hereby express their thanks for the cooperation of Donor Network West and all of the organ and tissue donors and their families, for giving the gift of life and the gift of knowledge, by their generous donation.

## Consent statement

All human subjects provided informed consent.

## Author contributions

Y.A.A., A.L.N., M.M., W.W.S, R.V.F., and T.C.W. conceived the project, and R.V.F. and T.C.W. acquired project funding. A.L.N, W.W.S, provided brain tissue samples. SS helped with data analysis. Y.A.A. generated reagents, performed lipidomics and proteomics experiments and data analysis, with help from S.S. A.L.N, and W.W.S, provided clinical data. S.K. contributed with sample preparations. Z.L., helped for proteomics data analysis. Y.A.A., R.V.F., and T.C.W. co-wrote the manuscript with input from all authors.

