## Supplemental figures for "Shared lipidome and proteome signatures of frontotemporal lobar degeneration and Alzheimer’s disease"

[5]Present address: Department of Pediatrics/Neurology at Baylor College of Medicine

**Running title**: Shared lipidome and proteome alterations in FTLD and AD

### Supplementary figure legends

**Supplementary fig. 1 | Lipidomic alterations in occipital cortex of FTLD compared with neurologically normal controls.** **A** PCA of lipidomic profiles from occipital cortex samples demonstrates separation between FTLD and control groups, indicating disease-associated lipid remodeling. **B** Quantification of major lipid classes in occipital cortex from control and FTLD subjects. Bars represent mean ± SEM, with individual data points shown. Significant differences between groups are indicated by asterisks (*$P < 0.05$, statistical test described in Methods).

**Supplementary fig. 2 | Alterations in BMP molecular species and ganglioside classes in FTLD frontal cortex.** **A** Fold-changes of individual BMP molecular species in FTLD relative to controls. **B** Fold-changes of major ganglioside subclasses in FTLD relative to controls. Data are shown as mean ± SD, with individual samples overlaid. Statistical significance was assessed using multiple unpaired two-tailed Mann-Whitney U tests. *$P < 0.05$; nd, not detected.

**Supplementary fig. 3 | Global lipidomic alterations in the occipital cortex of FTLD cases relative to controls.** **A** Distribution of log2 fold-changes for individual lipid species in FTLD relative to controls across lipid classes. Each point represents a single lipid species, illustrating the magnitude and direction of disease-associated lipid alterations. **B** Volcano plot showing differential abundance of lipid species in FTLD occipital cortex compared with controls. The x-axis represents log2-fold change (FTLD/Control), and the y-axis represents −log10(P value). Red and blue points indicate significantly increased and decreased lipid species, respectively, and gray points represent non-significant changes. Selected significantly altered lipid species are annotated. Dashed lines indicate fold-change and significance thresholds.

**Supplementary fig. 4 | Lipidomic alterations across FTLD subtypes in frontal cortex.** **A** Heatmap showing relative abundance of lipid classes across controls and FTLD subtypes **B–F** Volcano plots of differentially abundant lipid species in each FTLD subtype relative to controls, highlighting significantly increased and decreased lipids in FTLD-*GRN* (**B**), FTLD-*C9orf72* (**C**), FTLD-A-Sporadic (**D**), and FTLD-C-Sporadic (**E**) and FTLD-tau-PiD (**F**).

**Supplementary fig. 5 | Proteomic alterations in the occipital cortex of FTLD.** **A & B** PCA of proteomic profiles from frontal and occipital cortex comparing FTLD and control subjects. **C** Volcano plot showing differentially abundant proteins in FTLD occipital cortex relative to controls. **D** Pathway enrichment analysis of significantly altered proteins, highlighting dysregulated metabolic, lipid metabolic, and lysosomal-related pathways.

**Supplementary fig. 6 | Lipidomic and proteomic alterations in AD.** **A** Fold-changes of BMP molecular species in AD frontal cortex relative to controls. **B** Fold-changes of ganglioside classes in AD frontal cortex relative to controls. **C** Distribution of log2 fold-changes across lipid classes in AD occipital cortex. **D** Volcano plot of differentially abundant lipid species in AD occipital cortex. **E** Volcano plot of differentially abundant proteins in AD occipital cortex. **F** Pathway enrichment analysis of altered proteins. Statistical significance was assessed using multiple unpaired two-tailed Mann-Whitney U tests. *$P < 0.05$; nd, not detected.

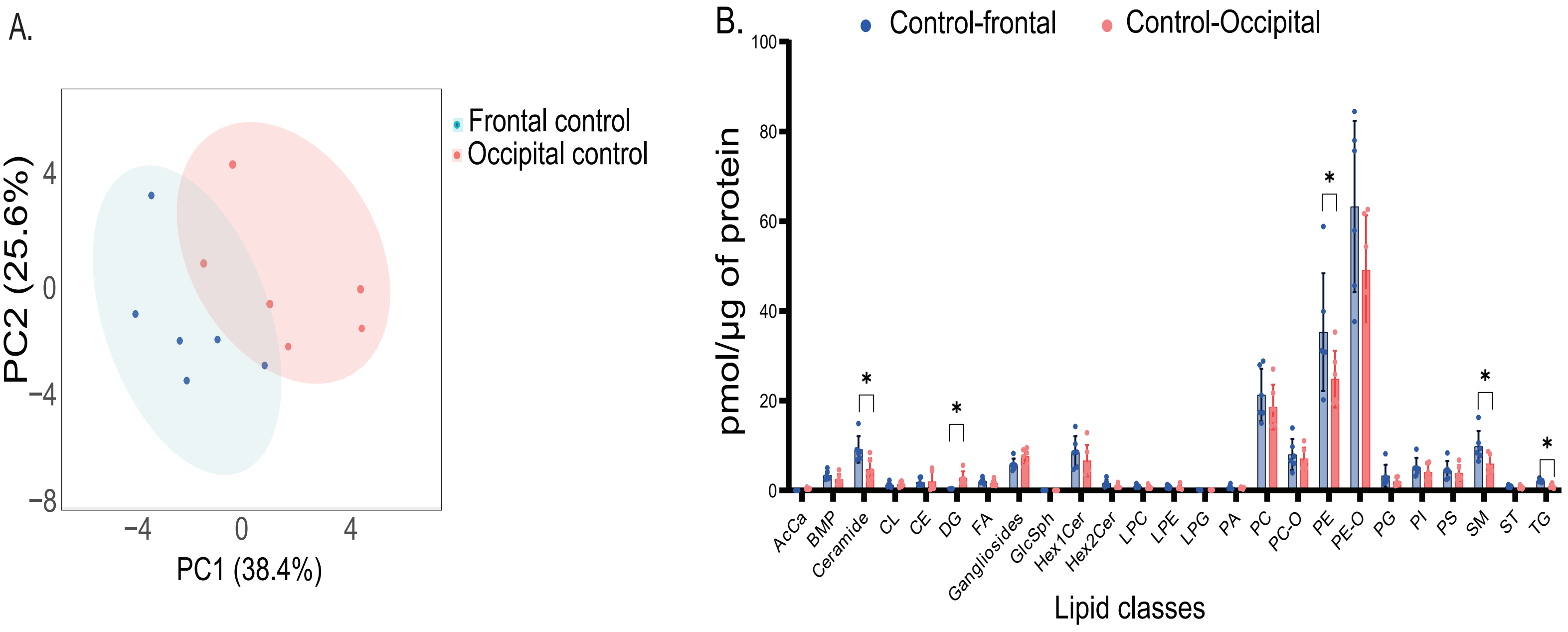
A.
Frontal control
Occipital control
PC2 (25.6%)
PC1 (38.4%)
4
0
−4
−8
−4
0
4
B.
Control-frontal
Control-Occipital
pmol/µg of protein
100
80
60
40
20
0
*
*
*
*
*
AcCa
BMP
Ceramide
CL
CE
DG
FA
Gangliosides
GlcSph
Hex1Cer
Hex2Cer
LPC
LPE
LPG
PA
PC
PC-O
PE
PE-O
PG
PI
PS
SM
ST
TG
Lipid classes

Supp figure 2

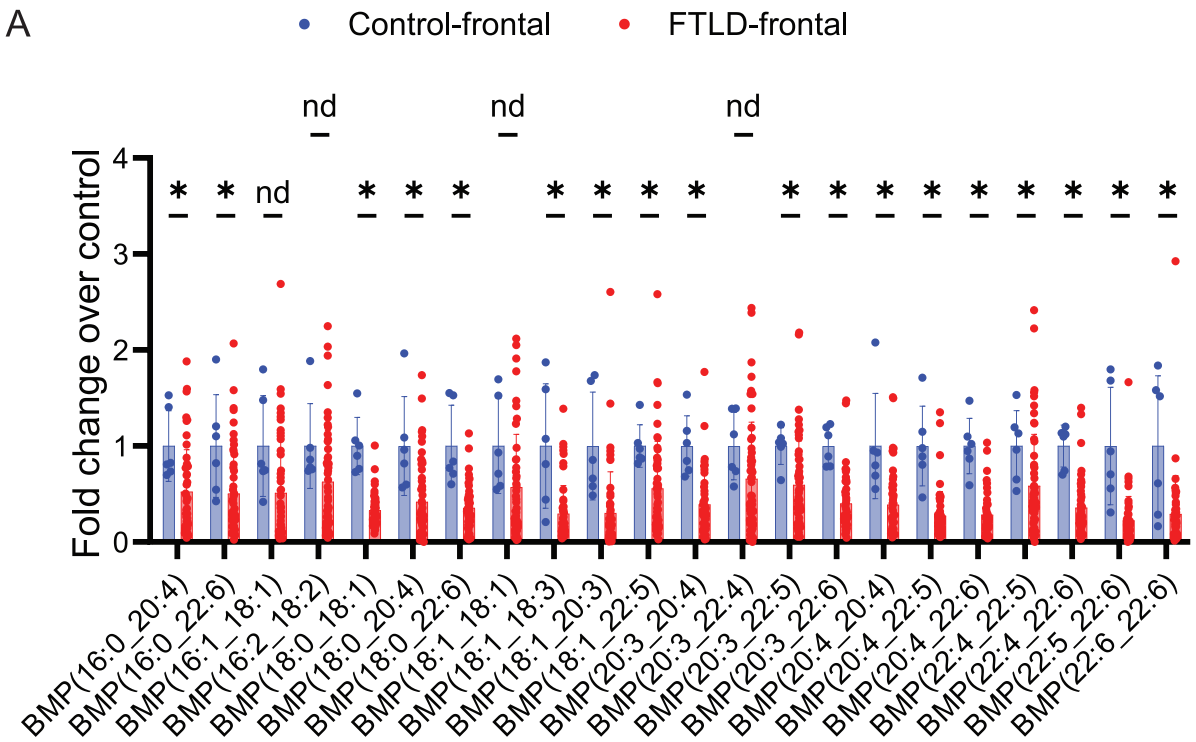


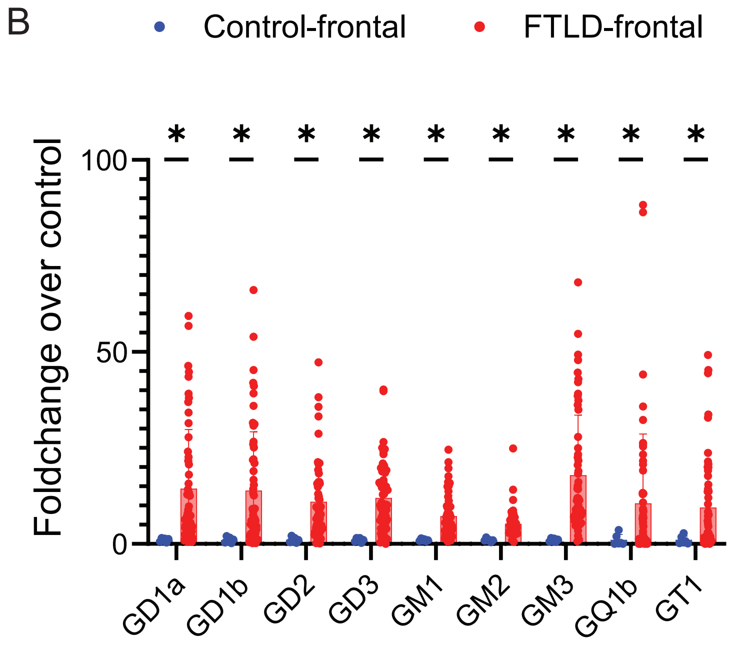

Supp figure 3

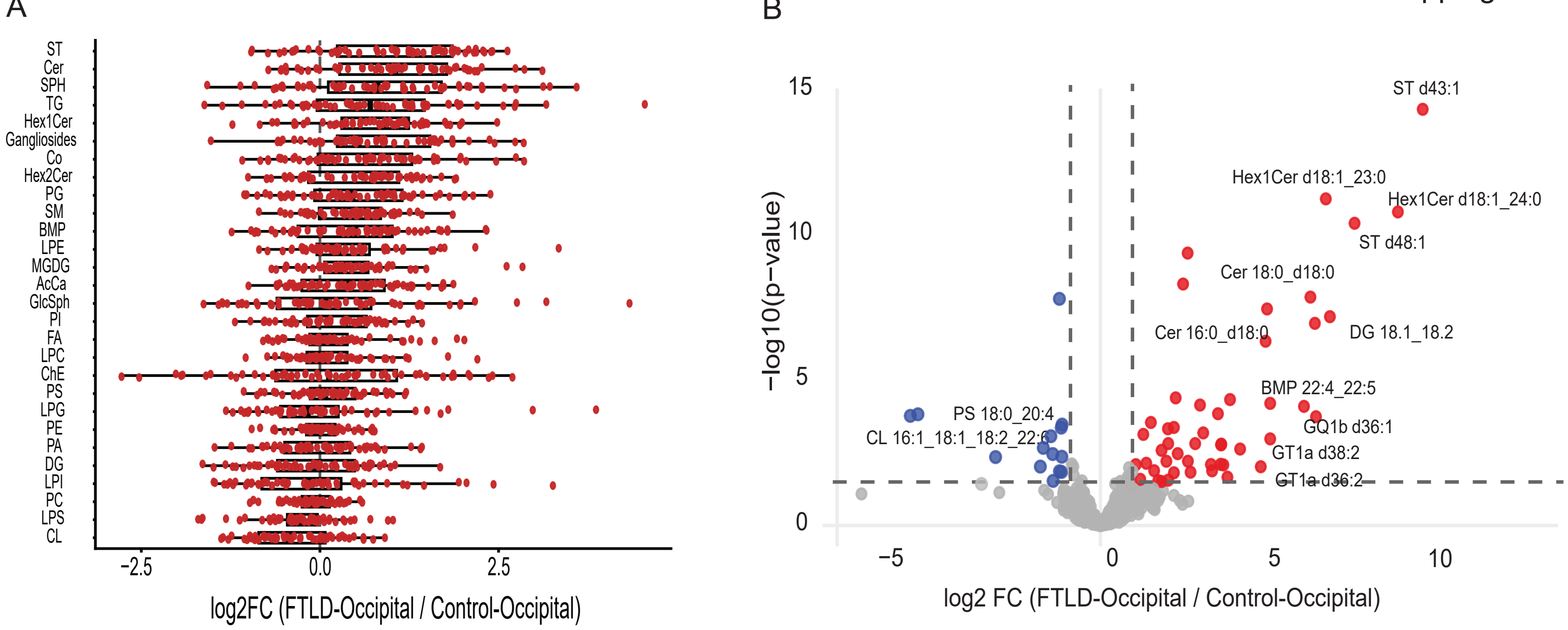

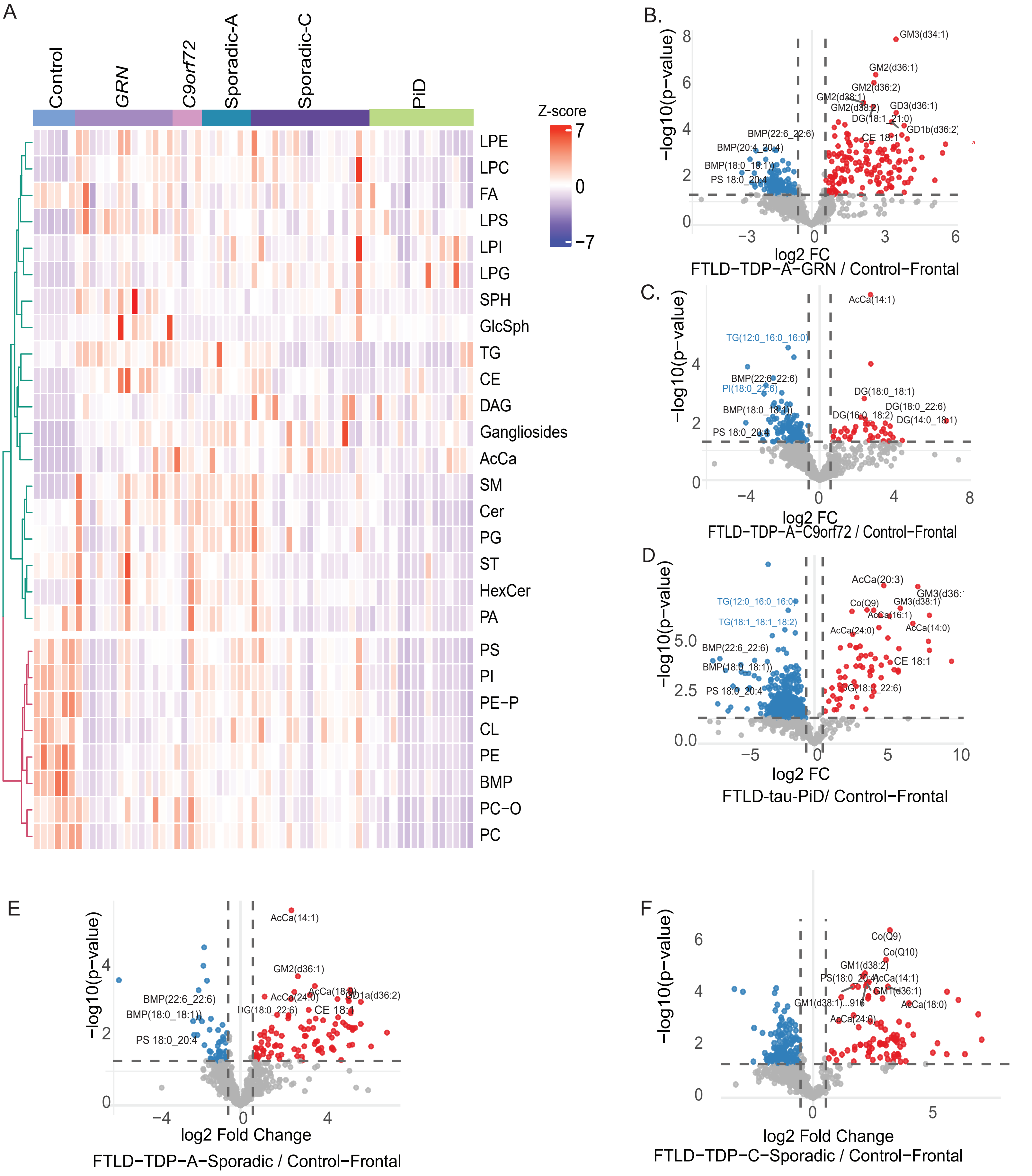
A
Control
GRN
C9orf72
Sporadic-A
Sporadic-C
PiD
Z-score
7
0
−7
LPE
LPC
FA
LPS
LPI
LPG
SPH
GlcSph
TG
CE
DAG
Gangliosides
AcCa
SM
Cer
PG
ST
HexCer
PA
PS
PI
PE-P
CL
PE
BMP
PC-O
PC
B.
−log10(p−value)
log2 FC
FTLD−TDP−A−GRN / Control−Frontal
GM3(d34:1)
GM2(d36:1)
GM2(d36:2)
GD3(d36:1)
DG(18:1_21:0)
CE 18:1
BMP(22:6_22:6)
BMP(20:4_20:4)
BMP(18:0_18:1)
PS 18:0_20:4
C.
−log10(p−value)
log2 FC
FTLD−TDP−A−C9orf72 / Control−Frontal
AcCa(14:1)
TG(12:0_16:0_16:0)
BMP(22:6_22:6)
PI(18:0_22:6)
BMP(18:0_18:1)
PS 18:0_20:4
DG(18:0_18:1)
DG(18:0_22:6)
DG(14:0_18:1)
D
−log10(p−value)
log2 FC
FTLD-tau-PiD/ Control−Frontal
AcCa(20:3)
TG(12:0_16:0_16:0)
Co(Q9)
GM3(d38:1)
AcCa(16:1)
TG(18:1_18:1_18:2)
AcCa(24:0)
AcCa(14:0)
BMP(22:6_22:6)
BMP(18:0_18:1)
CE 18:1
PS 18:0_20:4
E
−log10(p−value)
log2 Fold Change
FTLD−TDP−A−Sporadic / Control−Frontal
AcCa(14:1)
GM2(d36:1)
BMP(22:6_22:6)
AcCa(24:0)
AcCa(18:0)
BMP(18:0_18:1)
CE 18:1
PS 18:0_20:4
F
−log10(p−value)
log2 Fold Change
FTLD−TDP−C−Sporadic / Control−Frontal
Co(Q9)
Co(Q10)
GM1(d38:2)
AcCa(14:1)
GM1(d36:1)
AcCa(18:0)
AcCa(24:0)

Supp_figure 5

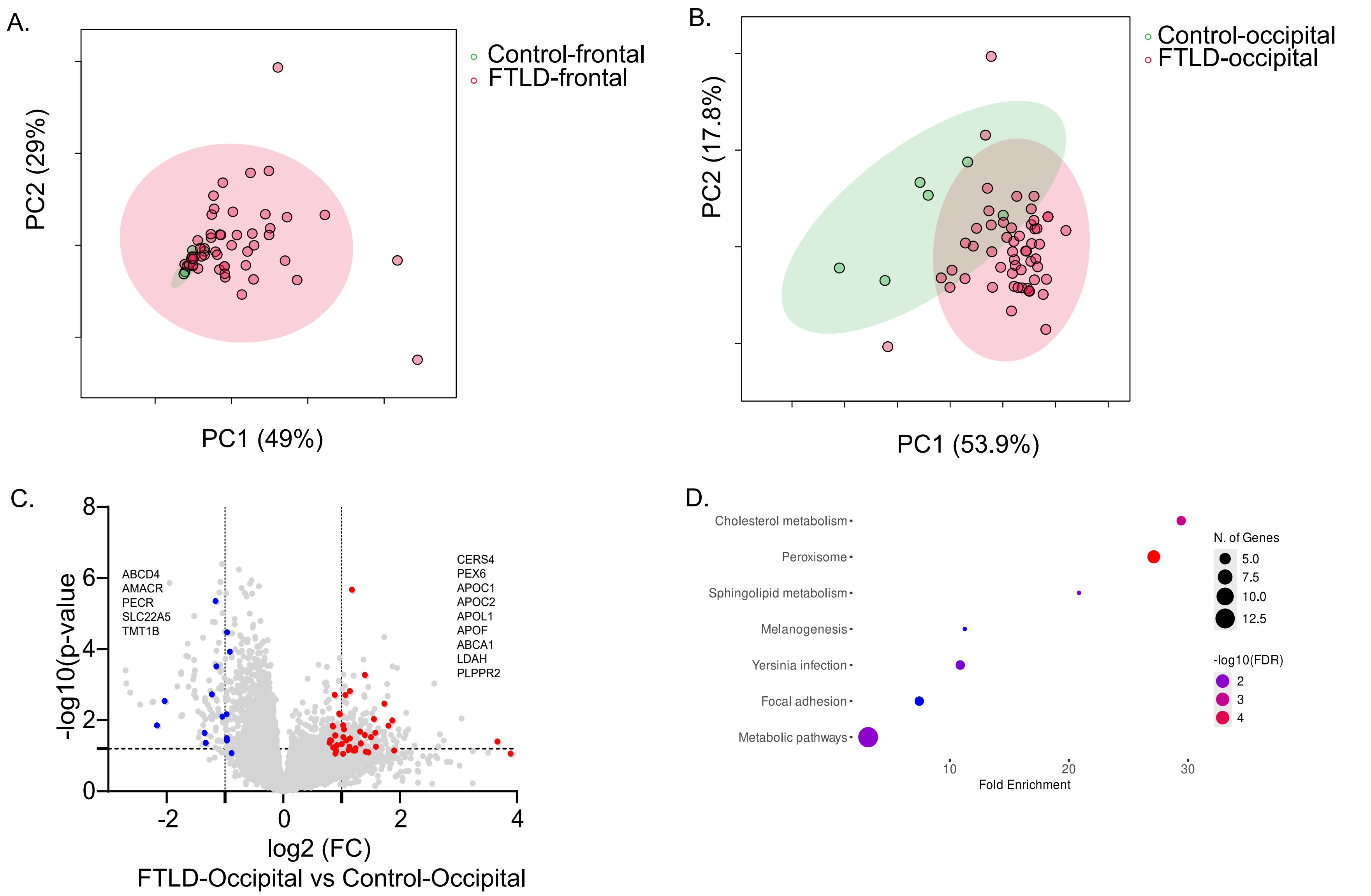

Supp_figure 6

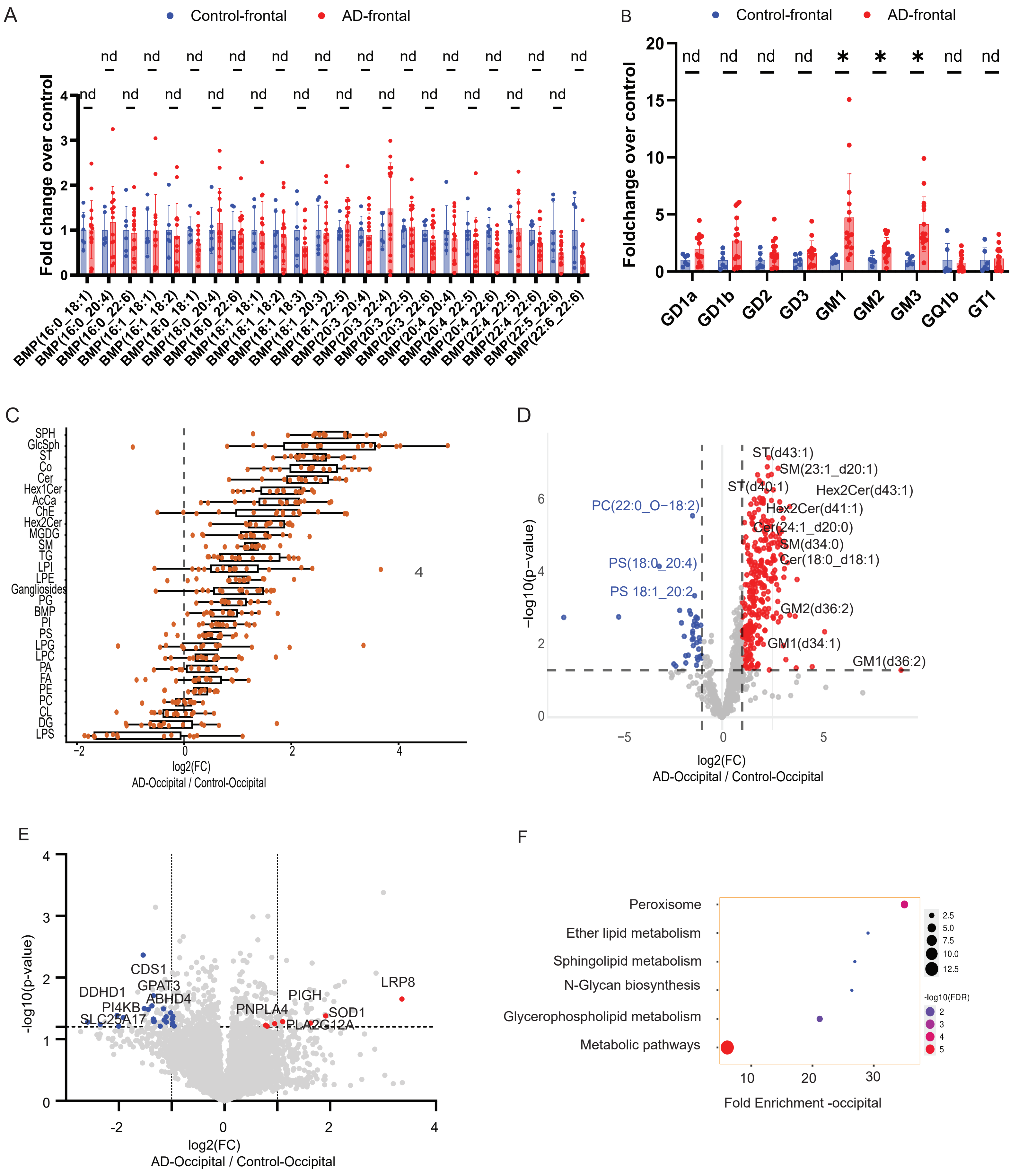
